# A low dose of simvastatin enhanced the therapeutic efficacy of mesenchymal stem cell (MSC) transplantation in skin wound healing in diabetic mice associated with increases in pAkt, SDF-1, and angiogenesis

**DOI:** 10.1101/763417

**Authors:** Supakanda Sukpat, Nipan Israsena, Jutamas Wongphoom, Praewphan Ingrungruanglert, Tao Ming Sim, Suthiluk Patumraj

## Abstract

**Purpose:** We aimed to determine the possible mechanisms of underlying the effects of low dose simvastatin on enhancing the therapeutic efficacy of MSC transplantation in diabetic wound healing.

**Methods:** Balb/c nude mice were divided into five groups:- control mice (CON), diabetic mice (DM), diabetic mice pretreated with low-dose simvastatin (DM+SIM), diabetic mice implanted with MSCs (DM+MSCs) and diabetic mice pretreated with low-dose simvastatin and implanted with MSCs (DM+MSCs+SIM). Seven days before wound induction, low dose simvastatin was orally administered to the DM+SIM and DM+MSCs+SIM groups. Eleven weeks after the induction of diabetes, all mice were given bilateral full-thickness excisional back skin wounds.

**Results:** By comparing the DM+MSCs+SIM and DM+MSCs groups, the results showed that on day 14; the wound closure (%WC) and capillary vascularity (%CV) in the DM+MSCs+SIM group were significantly increased compared to those in the DM+MSCs group. In addition, by using immunohistochemical techniques, it was also shown that the expression of SDF-1, a chemotactic factor regulating the migration of stem cells, in the DM+SIM+MSCs group was increased compared with that in the DM+MSCs group. Furthermore, using phospho-Akt (S473) Pan Specific DuoSet IC ELISA (R&D Systems, USA) kits, the increased tissue Akt levels were found in the DM+SIM+MSCs group but not in the other groups.

**Conclusions:** Our study suggests that a low dose of simvastatin enhanced the therapeutic efficacy of MSCs in diabetic wound healing, and this effect was associated with increases in pAkt levels, SDF-1 levels, and angiogenesis, and improved wound closure.

## Introduction

In diabetes, hyperglycemia-induced oxidative stress resulting from ROS production has been observed in several cell types [1]. Studies have shown that hyperglycemia-induced oxidative stress could lead to impaired wound healing via inducing a prolonged chronic inflammatory state. This prolonged inflammatory phase could in turn disturb collagen metabolism, resulting in poor blood supply, the reduced production of growth factors, and reduced angiogenesis, all of which contribute to delayed diabetic wound healing [2–4]. Nearby 2% to 3% of DM patients suffer from active foot ulcers [5–6]. Diabetic foot ulcers not only affect the physical health of patients, but they lead to 85% of major lower limb amputations in patients with DM. Furthermore, patients and society must bear a substantial financial burden resulting from the treatment and care of diabetic foot ulcers [7].

Currently, novel therapeutic interventions involving cellular therapies and tissue engineering approaches are being rapidly developed. In the context of diabetes research, accumulating evidence has indicated that mesenchymal stem cells (MSCs) could improve wound healing [8–10]. However, it has been suggested that the consequences of hyperglycemia-induced oxidative stress could cause problems because of decreased MSC paracrine signaling due to the induction of replicative senescence and apoptosis [11–12]. In other words, the therapeutic functions of MSCs could be impaired resulting in low capacity and poor quality in MSC transplantation for diabetic ulcer treatment.

A major problem in the use of MSCs is the transplantation environment [13]. For example, 90% of MSCs were shown to be apoptotic, which resulted in programmed cell death, within 3 days after transplantation into the heart for cardiomyocyte infarct repair [14–15]. In the last few years, the effects of inflammation induced immunosuppression has been taken into account in the use of rational design for the clinical use of MSCs. This indicates that, first, an optimal administration time point should be carefully selected according to the levels and ratios of different cytokines in the body during disease progression. Second, cytokine priming should be attempted to improve the therapeutic effect of MSCs. Third, the therapeutic efficacy of MSCs probably depends on the nature of different diseases because of the distinct inflammatory environments that are involved.

In addition, accumulating evidence has revealed that the therapeutic benefits of MSCs are largely dependent on their capacity to act as a *trophic factor* pool. After MSCs home into the damaged tissue sites for repair, they will closely interact with the local stimuli, such as inflammatory cytokines, the ligands of Toll-like receptors, and hypoxia condition, all of which stimulate MSCs to produce a large amount of growth factors that are involved in multiple functions during tissue regeneration [16–17]. An improved understanding of the molecular pathways involved in growth factor production would be very helpful for developing better strategies for MSC-based therapies.

In our previous study, by using daily oral feeding of simvastatin (0.25 mg/kg/day) in diabetic mouse model, the pleiotropic effects of simvastatin including reduced neutrophil infiltration, enhanced VEGF production, increased angiogenesis, and improved wound healing were demonstrated [18].

A number of studies have shown that simvastatin could reduce inflammatory infiltration, enhance VEGF production, increase the number of circulating endothelial progenitor cells (EPCs), increase angiogenesis, and improve wound healing in diabetic and nondiabetic models [19–23]. In addition, the immediate effects of simvastatin could improve the efficacy of MSCs, increased VEGF production and increased angiogenesis in hindlimb ischemia, a stroke model and several in vitro models [24–27].

By considering these previous findings, we hypothesized that the anti-oxidative stress and anti-inflammatory effects of low-dose simvastatin supplementation will provide the optimal microenvironment for improving the therapeutic efficacy of MSC transplantation in a diabetic wounds. By comparing groups of diabetic mice pretreated with low-dose simvastatin and implanted MSCs (DM+MSCs+SIM) and diabetic mice implanted MSCs (DM+MSCs), the results showed that on day 14; the wound closure (%WC) and capillary vascularity (%CV) in the DM+MSCs+SIM group were significantly increased when compared to those in the DM+MSCs group. In addition, by using immunohistochemical techniques, it was also shown that the expression of SDF-1, a chemotactic factor regulating the migration of stem cells, in the DM+SIM+MSCs group was increased compared with that in the DM+MSCs group. Furthermore, by using phospho-Akt (S473) Pan Specific DuoSet IC ELISA (R&D Systems, USA) kits, increased tissue Akt levels were found in the DM+SIM+MSCs group but not in the other groups. Our study suggests that a low dose of simvastatin enhanced the therapeutic efficacy of MSCs in diabetic wound healing, and this effect was associated with an increase in pAkt levels, SDF-1 levels, and angiogenesis, and improved wound closure.

## Materials and methods

### Animals

All procedures were conducted in accordance with guidelines for the use of experimental animals of The National Research Council of Thailand. The protocol for this study was approved by the Ethical Committee at the Faculty of Medicine, Chulalongkorn University, Bangkok, Thailand (approval ID:009/58). Male BALB/c nude mice (8 weeks old, body weight 20–23 grams) obtained from the National Laboratory Animal Center, Salaya Campus, Mahidol University, Thailand, were used in this study. The animals were housed one per cage, and maintained under controlled conditions (a pathogen-limited room where the temperature was 25±3°C with a 12-hour light and 12-hour dark cycle). Animals were provided free access to standard laboratory feed and sterilized water. The animals were anesthetized with 55 mg/kg sodium pentobarbital (Ceva Santé Animale, France) when required.

The animals were randomly divided into five groups, as follows: group 1 is the control group (CON; n=12); group 2 was the untreated diabetic group (DM; n=12); group 3 was the diabetic group that received daily oral treatment with simvastatin (0.25 mg/kg) (DM+SIM; n=12); group 4 was the diabetic group that received the implantation of 1×10^6^ MSCs (DM+MSCs; n=12); group 5 was the diabetic group that received both daily oral treatment with simvastatin (0.25 mg/kg) and the implantation of 1×10^6^ MSCs (DM+MSCs+SIM; n=12).

### Induction and evaluation of diabetes

The diabetic mice were induced by intraperitoneal injection of streptozotocin (STZ; Sigma Chemical Co., USA, 45 mg/kg BW, daily for 5 days). Streptozotocin was freshly dissolved in citrate buffer, pH 4.5 (Sigma Chemical Co., USA). The same volume of citrate buffer, pH 4.5, was injected via the same route into nondiabetic control animals. Glucose levels were measured by a glucometer in tail vein blood samples obtaineds 2 weeks after diabetic induction. Diabetic conditions were defined as a plasma glucose concentration equal to or greater than 200 mg/dL [28].

### Cell culture and preparation of mesenchymal stem cells (MSCs)

Human bone marrow mesenchymal stem cells were obtained from the Stem Cell and Cell Therapy Research Unit at Chulalongkorn University. Briefly, MSCs were isolated from human bone marrow and were cultured in Dulbecco’s modified Eagle’s medium (DMEM) containing 5% fetal bovine serum (FBS), 1x penicillin-streptomycin solution, and L-glutamine at 37°C in 5% CO_2_. The medium was exchanged by removing the medium from the flask every 3-days during culture. The phenotypes of the MSCs were positive for CD44, CD73, CD90 and CD105 as shown in Table 1.

**Table 1.**
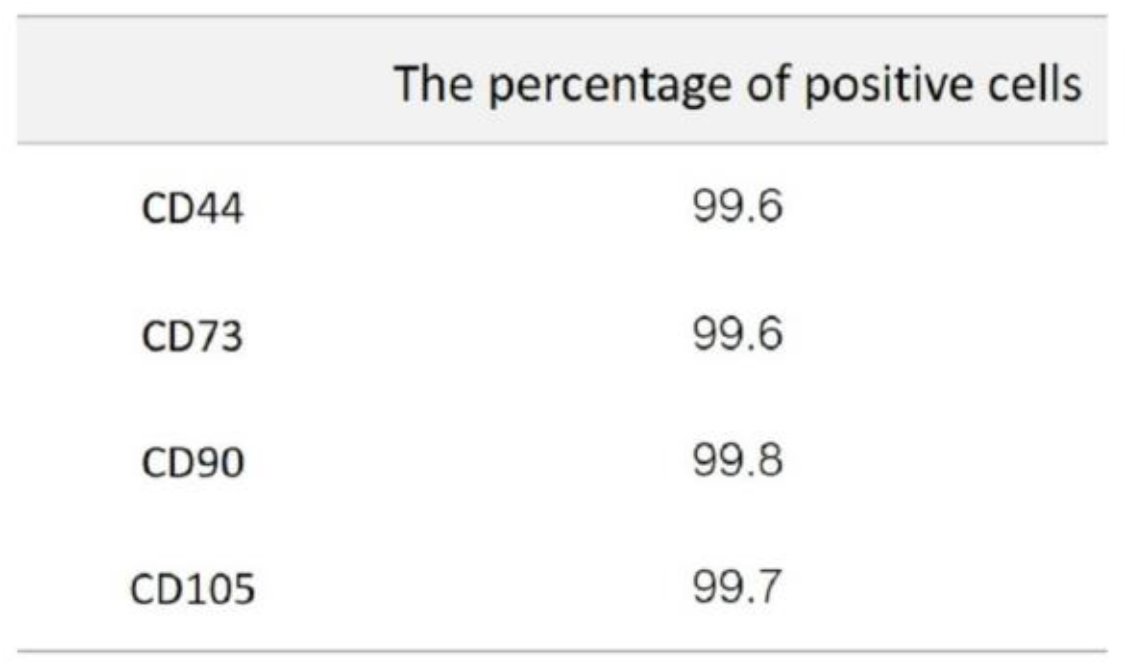
The percentage of positive MSCs. The percentage of positive MSCs were:-CD44; 99.6%, CD73; 99.6%, CD90; 99.8%, and CD105; 99.7%.

### Excisional skin wound model

In the 11th week after STZ induction, a cutaneous wound was generated. Briefly, after the induction of anesthesia by the intraperitoneal injection of sodium pentobarbital (55 mg/kg), a bilateral full-thickness excisional skin wound 0.6×0.6 cm^2^ in size was created on both sides of the midline of each mouse [18]. Fibrin gel (Shanghai RAAS-Blood Products Co. Ltd, China) or human mesenchymal stem cells were implanted in the wound bed. The wounds were then covered by square-shaped plastic splints and Tegaderm™ (3M Company, St. Paul, MN, USA) to decrease the possibility of contraction, thus ensuring that closure occurred via re-epithelialization during the wound healing process. Simvastatin supplements at a dosage of 0.25 mg/kg were given daily for 7 days before wound creation and continued to be given until the experimental end-point was reached [29–31].

### Measurement of wound closure

Photographs of wounds were taken by a digital camera on days 0, 7 and 14 post-wound creation. Areas of the wound were analyzed by digital image software analysis (Image-Pro Plus II 6.1; Media Cybernetics, Bethesda, MD), and the percentage of wound closure (%WC) was evaluated using the following equation: % WC = ((Area of original wound-Area of actual wound)/ Area of original Wound) x 100 [32].

### Measurement of capillary vascularity

After anesthetization, the jugular vein of each mouse was cannulated for injection of 0.1 ml of 5% FITC-labeled dextran (Sigma Chemical Co., USA). The wound area was then measured using confocal fluorescence microscopy at 100x magnification (Nikon Eclipse E800, Nikon, Japan), and the surrounding capillary vasculature was examined. From the fluorescent photographs of capillaries with diameters less than 15 µm, the percentage of capillary vascularity was calculated using Image-Pro II 6.1 software. The percentage of capillary vascularity was calculated using the following equation: %Capillary vascularity (%CV) = (Number of pixels within capillaries / Total number of pixels in the entire frame) x 100 [33].

### Measurement of neutrophil infiltration

Seven and 14 days after wound creation, the mice were sacrificed, and the tissues in the wound areas were harvested. The tissue specimens were fixed in 10% formaldehyde for 24 hours. Two-micrometer-thick sections were stained with hematoxylin-eosin (H&E). The number of neutrophils infiltrating in the wound areas was measured using light microscopy (Nikon Eclipse 50i, Nikon, Japan) at 400x magnification and Image-Pro II 6.1 software [34]. These results were confirmed by blind assessment.

### Determination of tissue VEGF level

Seven and 14 days after wound creation, tissue from the wound area was harvested and stored at −80°C for analysis of VEGF using an enzyme-linked immunosorbent assay (murine VEGF-specific ELISA kit (R&D Systems, USA) and a BCA protein assay kit (Thermo scientific, USA)). The amount of VEGF was expressed as picograms per milligram protein unit.

### Determination of SDF-1 level

The tissue SDF-1 level was determined using an immunohistochemical technique. Two-micrometer-thick paraffin-embedded sections were cut from the paraffin-embedded tissue. Heat-induced antigen retrieval was performed in citrate buffer (pH 9, Dako, USA). The sections were incubated with rabbit polyclonal SDF-1 antibody (sc-28876, Santa Cruz, USA, at 1:25 dilution). After washing in wash buffer, the sections were incubated with anti-rabbit secondary antibody (Dako, USA) and then with the DAB+ Substrate Chromogen System (Dako, USA) at room temperature. The sections were photographed at 400x magnification using a light microscope. Then, the images were analyzed using image software (Image-Pro Plus II 6.1). The results were confirmed by blind assessment.

### Determination of interleukin 6 (IL-6) levels and serine/threonine-specific protein kinase (Akt) activation

Homogenate supernatant samples were used for determining tissue IL-6 level and phospho-Akt (pAkt) levels. The levels were measured using commercially available IL-6 ELISA (R&D Systems, USA) and phospho-Akt (S473) Pan Specific DuoSet IC ELISA (R&D Systems, USA) kits, based on the manufacturer’s protocols [34–36]. The levels were expressed in picograms per milligram of protein.

### Determination of tissue malondialdehyde (MDA) levels

The supernatants of each tissue sample were used to analyze malondialdehyde levels (MDA; lipid peroxide parameter) by a TBARS assay kit (Cayman Chemical Co, USA). The MDA level was expressed as nM/mg of protein.

### Determination of MSC migration

Human MSC cells labeled with red fluorescent protein (RFP) were obtained from the Stem Cell and Cell Therapy Research Unit, Chulalongkorn University. Harvested cells (1×10^6^ cells in 60 µL fibrin gel) were implanted in the wound bed of each mouse.

To determine MSC migration, the mice were anesthetized with pentobarbital sodium and the jugular vein was cannulated for the injection of fluorescent tracers (FITC-labeled dextran). The dorsal skin surrounding the wound area was cut open and examined by confocal microscopy. After the injection of FITC-labeled dextran, confocal images of the microvasculature and MSCs-RFP cells in the wound areas were obtained at 10x magnification with absorption wavelengths and emission wavelengths of 492–549 nm and 518–565 nm, respectively.

### Statistical analysis

The results were expressed as the means ± standard errors of mean (SEM). The differences between groups were determined by one-way analysis of variance (ANOVA), followed by a least significant difference (LSD) post hoc test using SPSS software, version 21. Differences were considered significant at P < 0.05.

## Results

### Simvastatin produced the optimal microenvironment for improving the therapeutic efficacy of MSCs

In order to determine whether simvastatin could improve the therapeutic efficacy of MSCs as a result of its pleiotropic effects on nonhepatic effects tissue malondialdehyde levels (MDA), the number of local infiltrating neutrophils, and the levels of IL-6 in the wound area were evaluated as shown in Fig 1.

**Fig 1.**
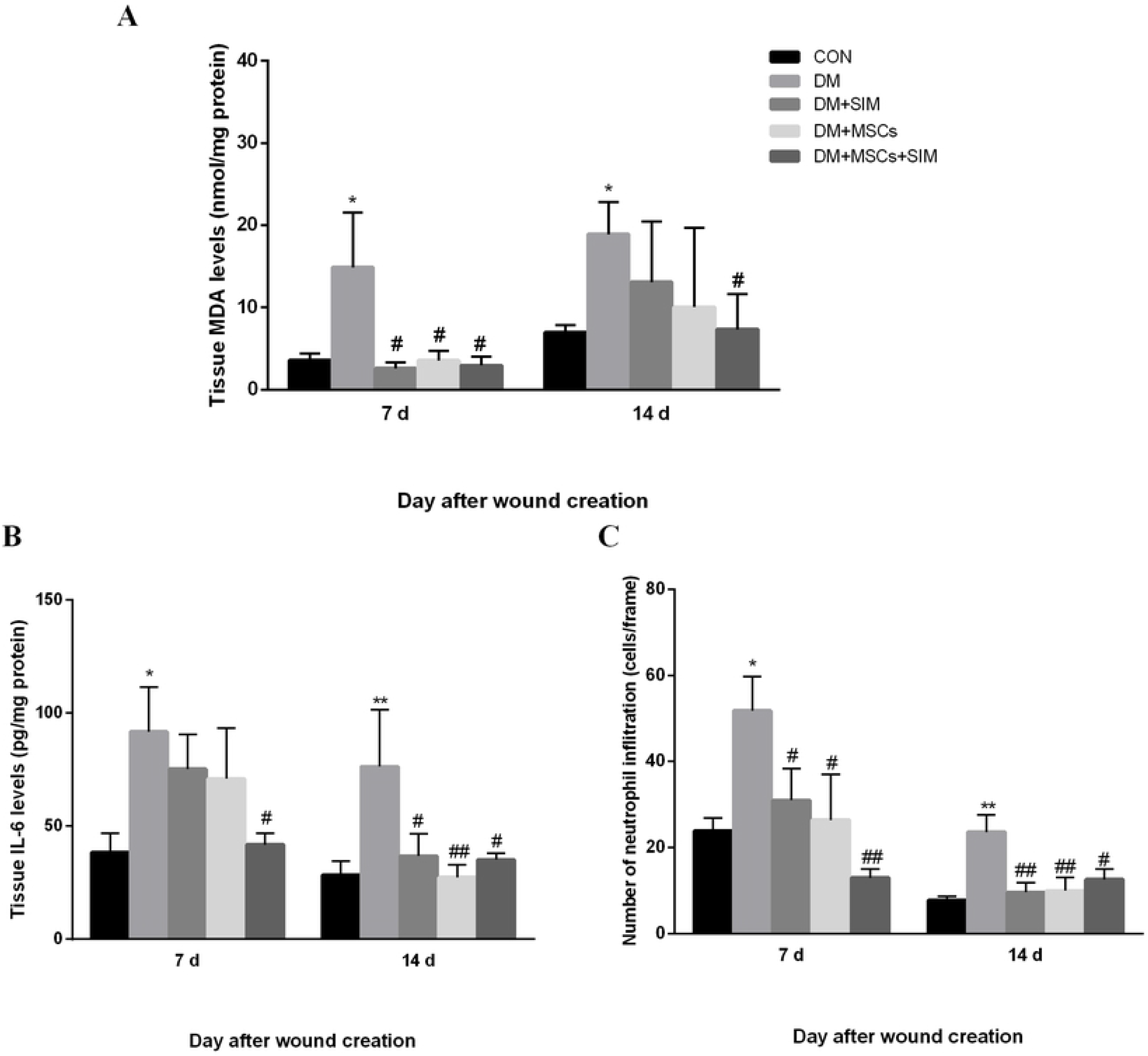
Effects of simvastatin on tissue malondialdehyde (MDA), local infiltrating neutrophils, and IL-6 in diabetic wounds. (A) Levels of tissue MDA were significant decreased on days 7 and 14 in DM+MSCs+SIM group. (B) Levels of tissue IL-6 on days 7 and 14 were were significant decreased on days 7 and 14 in DM+MSCs+SIM group. (C) The number of neutrophils infiltrating on days 7 and 14 were significant decreased in DM+MSCs+SIM group. The data are presented as the means ± SEM (n = 4-6). ANOVA, vs CON *p<0.05, **p<0.01; vs DM # p<0.05, ## p<0.01; ANOVA, vs CON *p<0.05; vs DM # p<0.05.

The increased in the tissue malondialdehyde levels (MDA, an ROS indicator), in the wound area; confirmed that oxidative stress contributed to poor diabetic wound healing (shown in Fig.1A). The consequent pathogenesis of wounds under diabetes induced oxidative stress, which was confirmed by high numbers of local infiltrating neutrophils, and high IL-6 levels, (Fig.1B, 1C), could be significantly attenuated by combined simvastatin and MSC treatment, as shown in the DM+MSCs+SIM group, however, our data showed that the effects of treatment were significant only in association with decreased anti-oxidant and IL-6 levels, but not in the increased numbers of local infiltrating neutrophils. In the present study, we found that treatments with both MSCs and simvastatin could reduce neutrophils infiltration in wounded diabetic mice on both days 7 and 14. On day 7 after wounding, the decrease in neutrophils infiltration in DM+MSCs+SIM group has showed a greater decrease than that in the DM+MSCs group, but this decrease was not significant. On 14-day after wounding, the results showed significantly decreased neutrophils infiltration in all treated groups, with no significant differences between the DM+MSCs+SIM and DM+MSCs groups. It might have been possible that the levels of neutrophils infiltration were the secondary effects after the anti-inflammatory phase, therefore, they did not reflect the significant improvement of MSC efficacy via the enhancement by simvastatin.

### Simvastatin enhanced the therapeutic efficacy of MSCs on wound angiogenesis

To determine whether simvastatin could enhance the efficacy of MSCs in inducing wound angiogenesis, the percentage of capillary vascularity (%CV) in the wound area was evaluated as shown in Fig 2.

**Fig 2.**
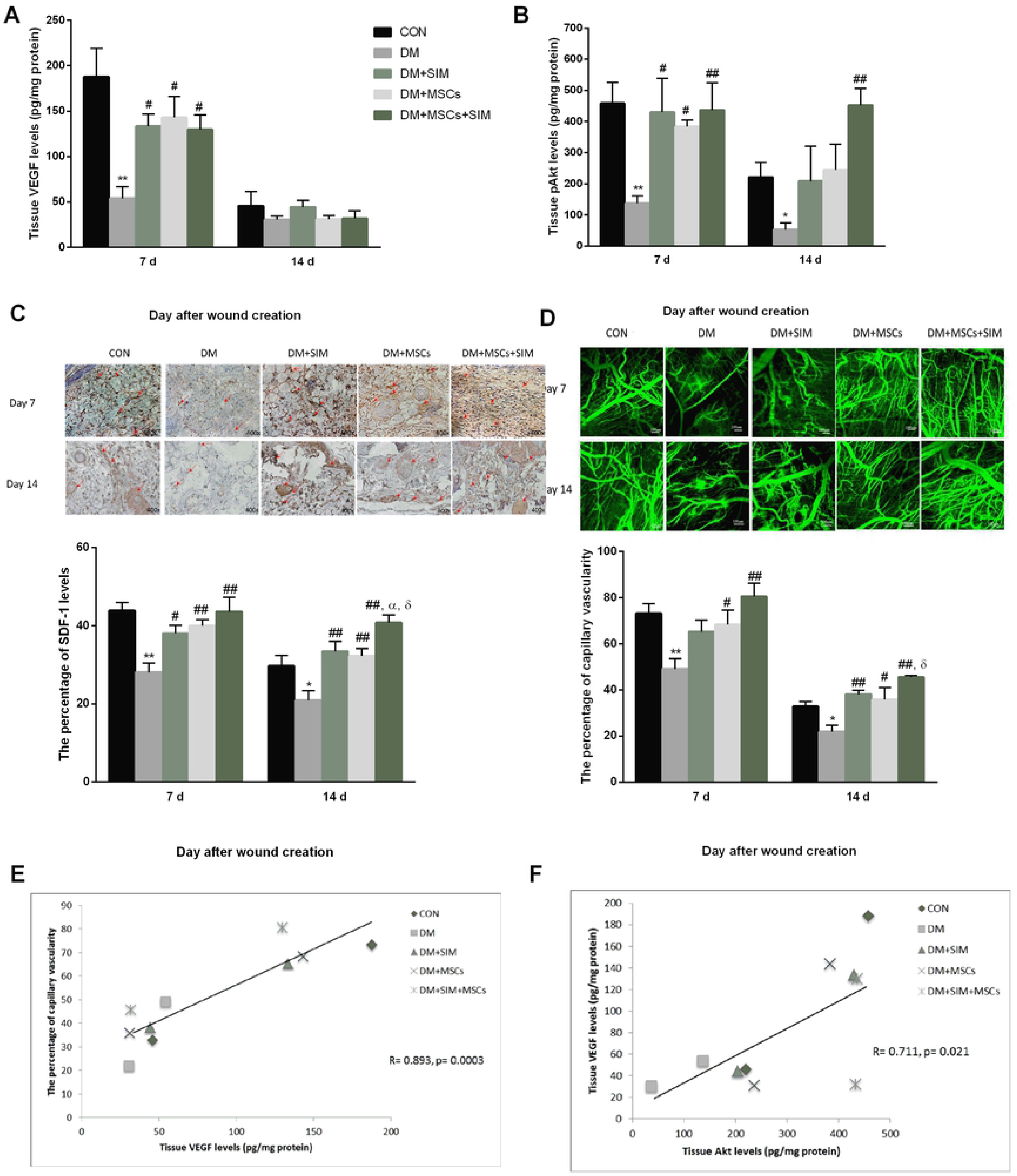
Effects of simvastatin on the therapeutic efficacy of MSCs on wound angiogenesis. (A) On day 7 after wound creation, VEGF levels were significantly higher in the DM+SIM, DM+MSCs, and DM+MSCs+SIM groups than in the DM group. (B) On day 7 and day 14, tissue pAkt were significantly higher in the DM+MSCs+SIM than that in the DM group. (C) On day 7 and day 14, the levels of SDF-1 were significantly higher in the DM+MSCs+SIM than that in the DM group. (D) From the fluorescent photographs of capillaries with diameter less than 15 µm, on the day 14, the %CVs of all treatment-groups were significantly higher than that in the DM group. (E-F) Pearson’s correlation analysis indicated close correlations between VEGF expression and %CV (R=0.893, p=0.0003); and between VEGF and the activation Akt/mTOR signaling (R=0.711, p=0.021). *indicates a significant difference vs CON p<0.05; vs DM, **p<0.01, # p<0.05, ## p<0.01; vs DM+MSCs, δ P<0.05. Abbreviations: VEGF, vascular endothelial growth factors; phospho-Akt, pAkt; %Capillary vascularity, %CV; CON, DM, DM+SIM, DM+MSCs and DM+MSCs+SIM

As a consequence of the pathogenesis of diabetes-induced oxidative stress, a reduction in the production of vascular endothelial growth factors (VEGF) and delayed wound healing was observed as shown in Fig. 2A. On day 7, VEGF levels were significantly lower in the DM group than in the control group. VEGF levels were significantly higher in the DM+SIM, DM+MSCs, and DM+MSCs+SIM groups than in the DM group (Fig 2A). However, on day 14, the results showed no significant difference among all groups.

The phosphatidylinositol 3-kinases (PI3Ks) and their downstream target AKT are considered crucial signal transduction enzymes that have been investigated extensively for their effects on cell proliferation, and angiogenesis in wound healing. Therefore, we conducted further investigation by using both ELISA and immunohistochemistry techniques to determine the effects of simvastatin on VEGF, pAkt and SDF-1 levels in the wound area (Fig 2A-C). On day 7 and day 14, pAkt and SDF-1 levels were significantly higher in the DM+MSCs+SIM than that in the DM group.

On day 14, a significant decreased %CV was observed in wounds in mice in the DM compared to that in wounds in the control mice group. Interestingly, on the day 14, the %CVs of all treatment-groups, including DM+SIM, DM+MSCs, and DM+MSCs+SIM, were significantly higher than that in the DM group (Fig 2D).

In addition, the Pearson’s correlation analysis indicated close correlations between VEGF expression and %CV (R=0.893, p=0.0003); and between VEGF and the activation Akt/mTOR signaling (R=0.711, p=0.021), as shown in Fig 2E-F.

In order to determine the effects of simvastatin on the local migration of human mesenchymal stem cells during the proliferation phase, the in vivo local migration of human mesenchymal stem cells labeled with red fluorescent protein (hMSCs-RFP) was observed by using confocal microscopy, (Fig.3A-D). In addition, the area of wound neovascularization (expressing green fluorescent FITC (Fig.3E) was also observed. By using Image Pro II software, the number of red fluorescent protein labeled MSCs homing in these existing vascular tubes was determined on the day 14 for the DM+MSCs and DM+MSCs+SIM groups (Fig 3F). The comparative results for the MSC-RFP vascular tubes seem to indicate the benefit effect of simvastatin on the enhancement of the effects of MSCs in wound angiogenesis.

**Fig. 3.**
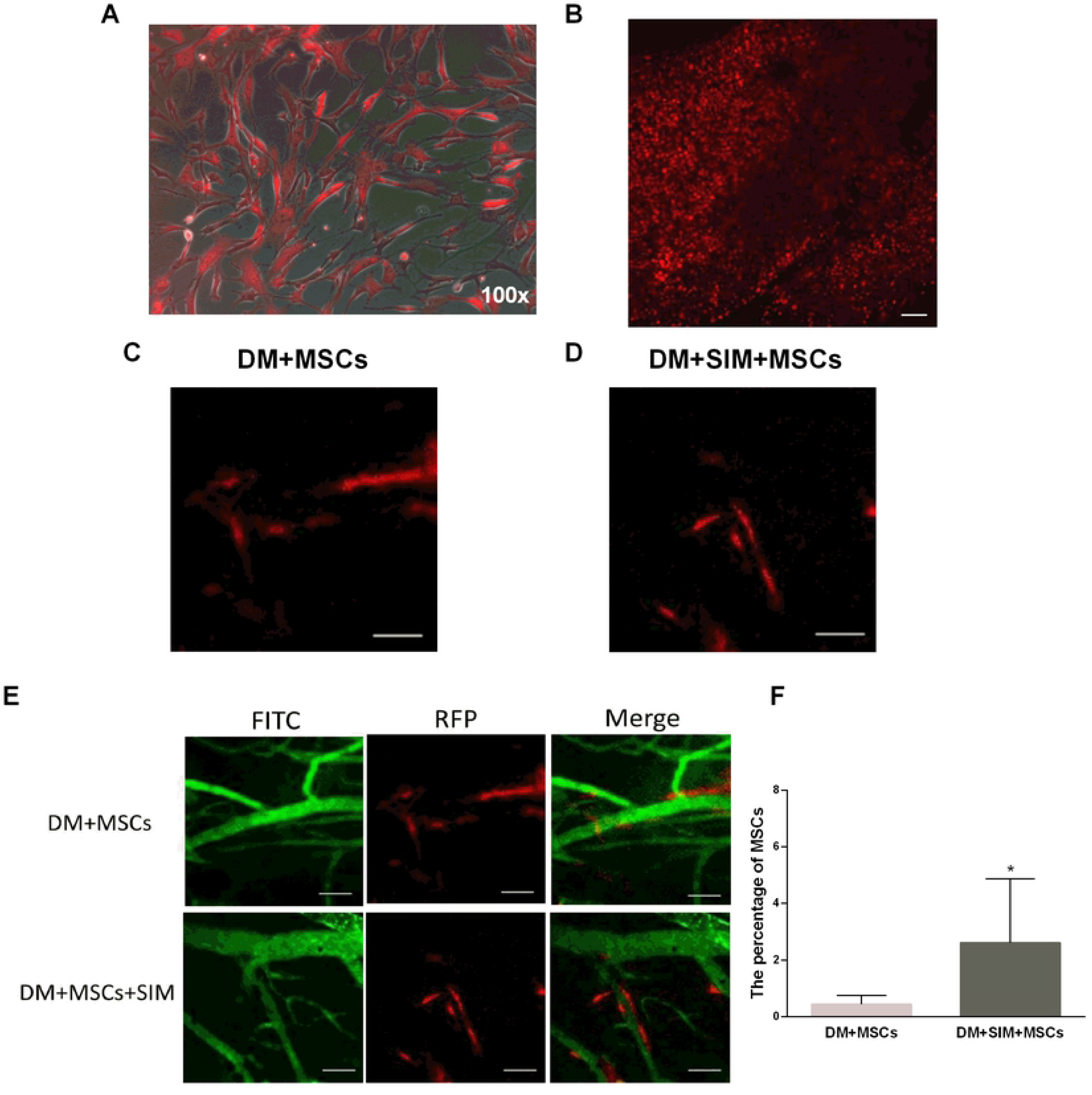
Effects of simvastatin on the local migration of human mesenchymal stem cells during the proliferation phase. (A) Microscopic photograph of human MSCs-RFP (red fluorescent protein stained) before the implantation. (B) Harvested cells (1×10^6^cells in 60 µL fibrin gel) were implanted to wound bed. (C) and (D) On 14-day after wound creation, the tube-like vessels were observed in DM+MSCs and DM+MSCs+SIM. (E) Representing confocal images of the FITC-labeled microvasculature (green) and MSCs-RFP cells (red) in wound areas. (F) The vascular tubes were significantly increased in the day 14 DM+MSCs+SIM than that in DM+MSCs group (*p<0.05).

### Simvastatin enhanced the therapeutic efficacy of MSCs in wound closure

On days 7 and 14 after wounding, the percentage of wound closure (%WC) was significantly lower in the DM group than in all other groups (Fig 4A). The %WC in DM+MSCs+SIM group was significantly higher than that in the DM+MSCs group on day 14.

**Fig 4.**
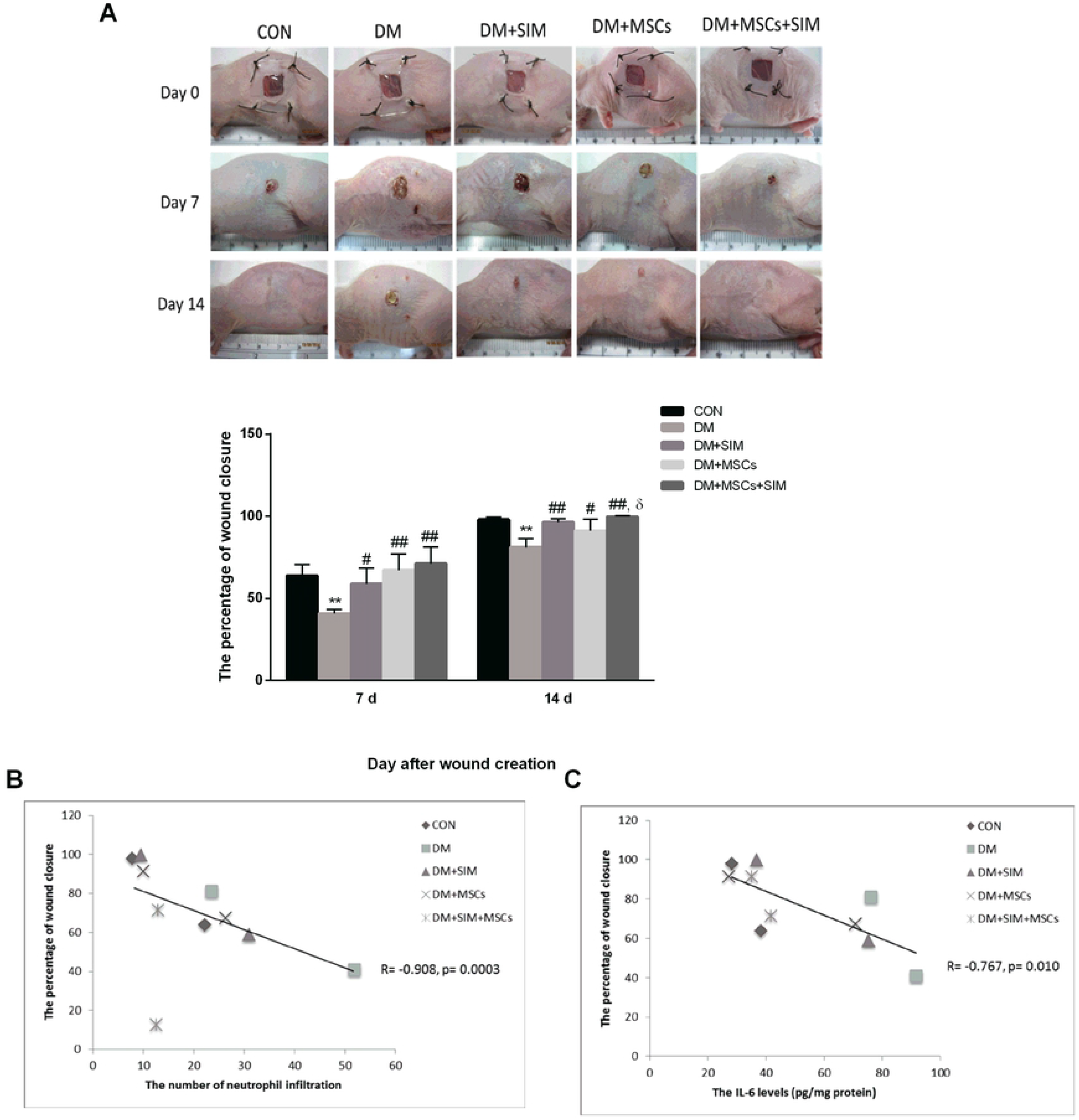
Effects of simvastatin on the therapeutic efficacy of MSCs on wound closure. (A) Representing photographs of the wound area in each group, and %WC on days 7 and 14 after wound creation were increases significantly in DM+MSCs+SIM than that in DM group. (B-C) Pearson’s correlation analysis indicated close correlations between neutrophils infiltration and %WC (R=0908, p=0.0003); and between IL-6 levels and %WC (R=0.767, p=0.01). Presented as the means ± SEM. (at day 7 and day 14, n=5-6 in each group). ANOVA, vs CON *P<0.01, vs DM # P<0.05, ## P < 0.01, vs DM+MSCs δ P < 0.05. Abbreviations: %WC, percent of wound closure

In addition, by using Pearson’s correlation analysis, the findings were also shown to confiorm the closed correlation between the expected factors representing the pleiotropic effects of simvastatin (number of neutrophils infiltration and IL-6 levels) and the therapeutic efficacy of MSCs on wound closure (%WC) (Fig. 4B; R=0.908, p=0.003) (Fig. 4C; R=0.767, p=0.010).

## Discussion

Diabetic wounds are characterized by the reduced production of growth factors, poor circulation and impaired angiogenesis, all of which contribute to poor wound healing. Our results agree with those of other studies that reported the excessive generation of reactive oxygen species (ROS) in diabetic wounds. Increased levels of tissue MDA, the ROS indicator, were found in the wound area, confirming that oxidative stress contributes to poor diabetic wound healing. The pathogenesis underlining diabetes-induced oxidative stress delayed wound healing involved high numbers of local infiltrating neutrophils, high IL-6 levels, and a reduction in the production of growth factors, such as platelet-derived growth factor (PDGF), transforming growth factor-β (TGF-β) and VEGF [37–38].

In the present study, we aimed to identify the therapeutic agent that could reduce the effects of contributing factors and improve diabetic wound healing. In the context of diabetes research, accumulating evidence has indicated that mesenchymal stem cells (MSCs) could improve wound healing and that the anti-oxidative stress and anti-inflammatory effects of low-dose simvastatin supplementation help to provide the optimal microenvironment for improving the therapeutic efficacy of MSC transplantation in a diabetic wound area.

Our findings have demonstrated that a low dose of simvastatin could enhance the therapeutic efficacy of MSCs in diabetic wound mice model. The benefitial effects of simvastatin combined with MSC treatment revealed by the acceleration of both wound closure and angiogenesis, which were increased almost 9% and 9.8% respectively compared to that in MSC-treated cells.

The results are consistent with the observation that paracrine factors play a key role in the contribution of MSC to wound healing, however, the limitation in the use of MSCs at injury sites, particularly those in a diabetic microenvironment, might be discouraging. In combination treatment with simvastatin, the anti-oxidant and anti-inflammatory effects of simvastatin could improve the therapeutic efficacy of MSCs in association with an increase in pAkt levels, SDF-1 levels, and improved angiogenesis.

Based on our findings, the massive decrease in neutrophils infiltration in wounds was observed in all treatment groups (DM+SIM, DM+MSCs and DM+MSCs+SIM groups). Previous studies suggested that MSCs could decrease neutrophil infiltration and inhibit neutrophil oxidative metabolism by upregulating the extracellular superoxide dismutase levels and reducing super-oxide anion concentrations and neutrophil extracellular trap formation [39].

Our results also showed that simvastatin could reduce neutrophil infiltration in diabetic wounds as well. The effects of simvastatin on reduction of extent of neutrophil infiltration were confirmed by studies performed in a skeletal muscle reperfusion injury model and a wound model. The stabilizing effect of simvastatin upon eNOS could enhance NO production during the early phase of reperfusion [40–42].

(Cowled et al. 2007, Karadeniz Cakmak et al. 2009)Similarly, the present finding demonstrated that simvastatin was able to reduce neutrophil infiltration in diabetic wounds, which might be explained by such mechanisms. The negative effects of statins on proinflammatory gene regulation results in the reductions of adhesive molecules, chemokines, cytokines, MMPs, and chemokine-receptor expressions. In addition, statins inhibit the pathway of inflammatory downregulated responses, which results in increased eNOS mRNA stability and NO production. Moreover, we believed that the anti-inflammatory effects of simvastatin may result in the optimization of the MSC-microenvironment, therefore, it wound be helpful to enhance the beneficial paracrine effects of BM-MSCs on wound healing.

Based on the confocal microscopy results, it was shown that 14 day after wounding, the percentage of capillary vascularity in the treatment groups (DM+SIM, DM+MSCs, and DM+MSCs+SIM) was significantly higher than that in the DM group. Moreover the findings showed that the percentage of capillary vascularity (%CV) in the group treated with combined simvastatin and MSC implantation significantly higher than that in the group that recieved MSC only. The formation of new blood vessels is considered to be one of the essential physiological processes in wound healing. The new blood vessels will increrase blood circulation and blood supply to the newly formed granulation tissue, and ensure keratinocytes survival, and re-epithelialization [43–44]. This observation is consistent with those in other reports of the therapeutic functions of MSCs in wound healing. In which neovascular network formation leads to a fast healing process [32–33]. Another aspect to take into account in term of the quality of wound angiogenesis is that on day 7 after wounding, increase in the level of vascular endothelial growth factors (VEGF) was observed in all treatment groups. VEGF is one of the key bioactive molecules implicated in the wound healing process that has been reported to possibly enhance endothelial cell proliferation and neovascularization, which is a critical stage in tissue regeneration by MSCs [47–48]. In the diabetic mouse model used in this study, the combination of simvastatin and MSCs was shown to increase the expression of SDF-1, which was demonstrated to have strong correlation with the Akt levels. Our results show that in the 14-day combined MSCs and simvastatin treatment group, the levels of SDF-1 expression increased almost 26% compared to that of the 14-day MSCs group, and tissue pAkt levels were increased compared with those in the DM group. The study performed by Zhao Y et al, 2016 [48] in ischemic stroke rat model confirmed a similar benefit effect of simvastatin and MSCs, and their results explored in greater details the increased expression of SDF-1 and VEGF associated with the activation of Akt/mTOR signaling pathway [48]. Phosphatidylinositol 3-kinases (PI3Ks) and their downstream target AKT are considered crucial signal transduction enzymes that have been investigated extensively for their roles in cell proliferation, cell transformation, paracrine function and angiogenesis [49–51]. Therefore, it is notable that the cytokine paracrine functions of BM-MSCs were enhanced by simvastatin leading to increased angiogenesis associated with PI3K/AKT pathway.

In addition, we investigated the contributions of SDF-1 to the recruitment of BM-MSCs to the wound area, and to the promotions of wound repair and neovascularization by labeling BM-MScs with red fluorescent protein (RFP) and then implanted them into the wound area with fibrin gel as described above. Our results show that on day 14 after wounding, the formation of vascular tubes in both the MSCs and combined MSCs and simvastatin groups was observed. The results of the present study together with the correlation analysis, imply that simvastatin may increase the migration of MSCs and the formation of vasculature in diabetic wounds in association with its significant correlation with the increase in SDF-1 levels and the activation of PI3K/Akt pathway.

Recent studies have indicated that, MSCs promoted the formation of vessel-like structures and the differentiation of tissue perivasculature using co-cultured endothelial cells with MSCs model [46, 52–53]. MSCs surrounding endothelial cells formed cord-like structures and expressed markers of pericytes such as alpha smooth muscle actin (α-SMA) and neuron-glial 2 (NG2), which are known to stabilize newly formed prevascular networks. (Verseijden etal. 2010, Rao etal. 2012, Pil etal.2015) Interestingly, simvastatin also enhanced MSCs migration and promoted MSCs-induced tube formation via acting on the SDF-1/CXCR4 signaling axis [54–55].

## Conclusion

These results may be the first in vivo evidence showing the synergistic effects of simvastatin and MSCs on the enhancement of angiogenesis in diabetic wounds associated with the PI3K/Akt pathway.

The proposed mechanism (Fig. 5) shows how low-dose simvastatin might enhance MSC efficacy in diabetic wounds. Briefly, simvastatin appears to enhance the effects of MSC treatment through pleiotropic mechanisms, including its effects on the increase in anti-oxidant, anti-inflammatory, and pro-angiogenic factors. The decreases in the inflammatory microenvironment are characterized by the extensive neutrophil infiltration and IL-6 involvement. Improved angiogenesis is reflected by increased tissue VEGF, SDF-1, and pAkt levels and thus by increased capillary vascularity. Finally, simvastatin enhanced the efficacy of MSCs in diabetic wound healing. The results of the present study indicated that the combined treatment with simvastatin and MSCs could reduce neutrophil infiltration, improve angiogenesis and increase VEGF levels, all of which could promote a better outcome for wound closure in a diabetic mouse model.

**Fig 5.**
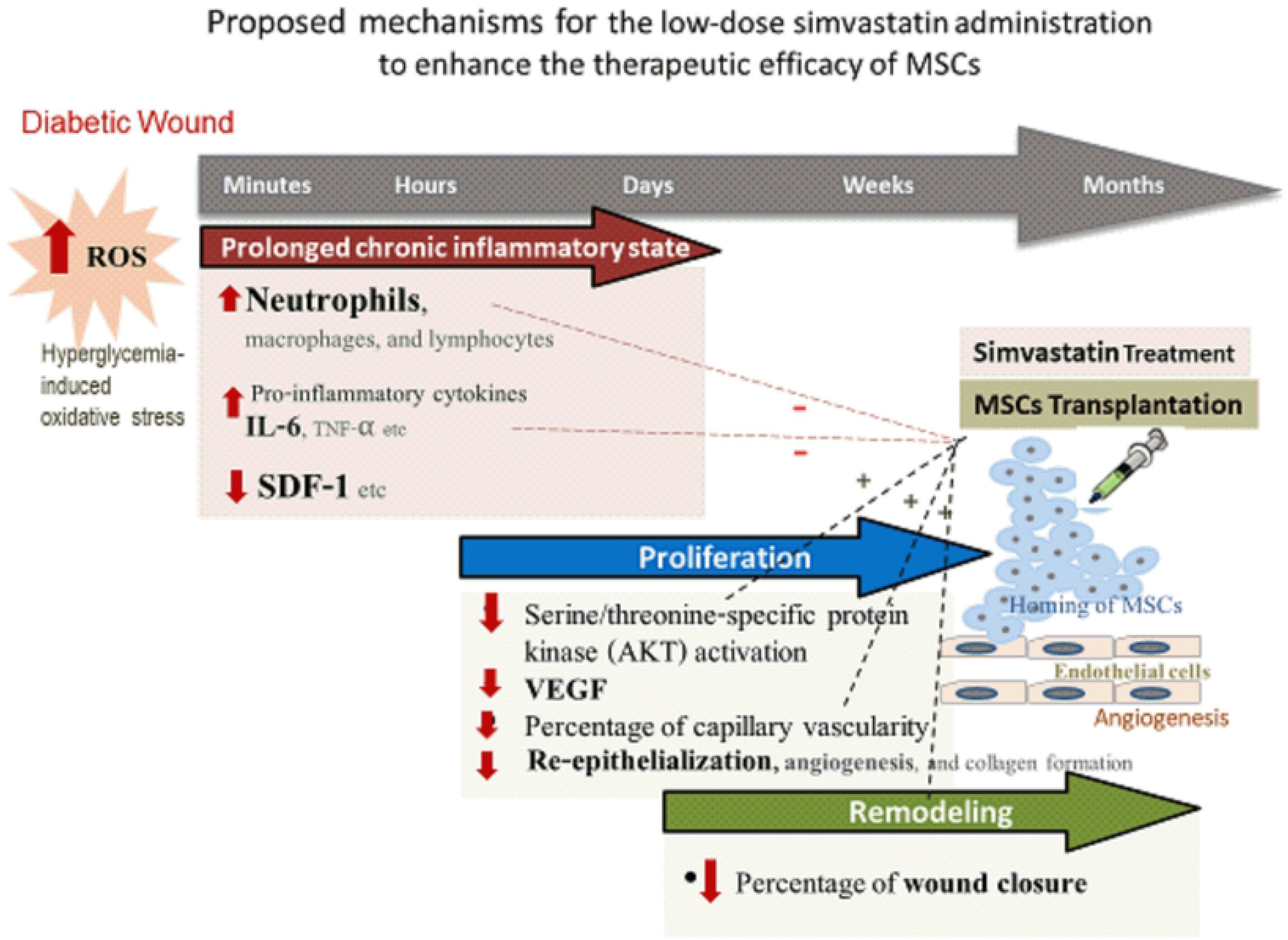
The proposed mechanism of how low-dose simvastatin enhanced MSCs efficacy in diabetic wounds.

## Acknowledgements

We thank Center for Excellence in Microcirculation, Department of Physiology, Faculty of Medicine, Chulalongkorn University, Bangkok, Thailand. The authors have no conflicts of interest to report.

## Supporting information

S1 Fig. This is the S1 Fig Title. This is the S1 Fig legend.

S2 Fig. This is the S2 Fig Title. This is the S2 Fig legend.

S3 Fig. This is the S3 Fig Title. This is the S3 Fig legend.

S4 Fig. This is the S4 Fig Title. This is the S4 Fig legend.

S5 Fig. This is the S5 Fig Title. This is the S5 Fig legend.

## Author Contributions

1. **Conceptualization:** SS NI SP.
2. **Data curation:** SS TMS SP.
3. **Formal analysis:** SS NI SP.
4. **Funding acquisition:** NI SP.
5. **Investigation:** SS JW TMS SP.
6. **Methodology:** SS NI JW TMS SP.
7. **Project administration:** SS NI SP.
8. **Supervision:** NI SP.
9. **Validation:** SS NI SP.
10. **Visualization:** SS TMS NI SP.
11. **Writing – original draft:** SS NI SP.

